# Characterization of Components of Resistance to Corn Stunt Disease

**DOI:** 10.1101/2020.05.28.120972

**Authors:** José D. Oleszczuk, María I. Catalano, Lucía Dalaisón, Julio A. Di Rienzo, María P. Giménez Pecci, Pablo D. Carpane

**Author notes:** corresponding author. P.D. Carpane.

## Abstract

Considering the occasional but increasing presence of corn stunt disease in the subtropical region of Argentina, the objective of this research was to devise an effective strategy to screen disease-resistant genotypes in the absence of high and constant natural pressures. To do so, the presence of antixenosis and antibiosis as components of resistance to vector *Dalbulus maidis* (DeLong 1923) as well as resistance to the pathogen *Spiroplasma kunkelii* (Whitcomb et al. 1986) under artificial inoculation conditions were investigated in four widely-distributed maize hybrids in Argentina. The hybrids shown differences in the levels of resistance and target organisms (either the insect vector or the pathogen). Antixenosis and antibiosis to *D. maidis* were observed in DK72-10. Resistance of DK79-10 to *S. kunkelii* was evidenced by a delayed onset of symptoms, and DKB390 was antixenotic to *D. maidis* and highly resistant to *S. kunkelii*. A good association was found between symptom severity and yield, but not between symptom severity and accumulation of pathogen *S. kunkelii*. In conclusion, the proposed methodology was efficacious and can aid the screening of resistant genotypes in breeding programs to reduce the impact of corn stunt disease, ensuring that hybrids with good resistance level will be planted by farmers whenever disease occurs.

## Introduction

Corn stunt is one of the most significant diseases affecting maize crop in the Americas, because of its high prevalence and its potential to cause yield losses in endemic areas (Bajet and Renfro 1989; Bradfute et al. 1981; Oliveira et al. 1998). Following its initial detection (Alstatt 1945; Frazier 1945), its prevalence has increased in the Americas (Bradfute *et al*. 1981; Hruska *et al*. 1996; Giménez Pecci *et al*. 2002; 2005). Corn stunt disease was first confirmed in the subtropical region of Argentina during the 1990/91 crop season (Lenardón et al. 1993). High disease prevalence was later reported in this region in certain crop seasons (Giménez Pecci et al. 2002; Virla et al. 2003), and isolated symptomatic plants may be found occasionally in temperate areas of Argentina (Carloni 2010; Carloni et al. 2013; Giménez Pecci et al. 2002; 2003; 2005).

The mollicute *Spiroplasma kunkelii* (Whitcomb et al. 1986), known as Corn Stunt Spiroplasma (CSS), is the pathogen most commonly associated with corn stunt disease in Argentina. This mollicute is transmitted by leafhoppers, being *Dalbulus maidis* (DeLong 1923) the only vector species identified in field-collected samples in Argentina (Giménez Pecci et al. 2003; Virla et al. 1990; Paradell et al. 2001), although *Exitianus obscurinervis* (Stal 1859) (Hemiptera: Cicadellidae) was proven to be a vector species in experimental conditions (Carloni et al. 2011). The epidemiological significance of *D. maidis* lies in its high prevalence (Nault 1990), high transmission efficiency of the pathogen *S. kunkelii*, with no reduced longevity (Alivizatos and Markham 1986; Nault 1990), and the persistent-propagative transmission mode of the pathogen, so insects acquiring the pathogen remain inoculative throughout their lifespan (Nault 1980).

Symptoms of corn stunt disease appear three to five weeks after inoculation and typically include chlorotic stripes that appear near the leaf base (Nault 1980), extending further towards the leaf tips (Kunkel 1946) and may cover the entire leaf, as well as younger leaves. In other cases, leaves may exhibit reddening (Oliveira et al. 1998) or deformations (“cuts”) in the margins. Plants may have shortened internodes, which gives its name to stunting, and in some cases ear proliferation (Nault 1980). Symptoms are usually more severe towards the upper part of the plant and may impair the development of reproductive structures of the plants (Carpane et al. 2006; Nault 1980). Reduced yield resulting from corn stunt disease is directly related to symptom severity and the accumulation of the pathogen *S. kunkelii* (Virla et al. 2004), which are both highest if *S. kunkelii* is inoculated at early growth stages (Massola Junior et al. 1999b; Scott et al. 1977). In this situation, reduced yield may be high, ranging from 12 to 100% (Hruska and Gomez Peralta 1997; Massola Junior et al. 1999b; Oliveira et al. 2003; Scott et al. 1977).

One of the most convenient alternatives to reduce yield loss resulting from diseases is the use of resistant crops (Hogenboom 1993). The resistance of maize genotypes to *S. kunkelii* might remain stable over time and across regions due to the low genomic variation of this pathogen (Carpane et al. 2013). Field-resistant genotypes have in fact been obtained in areas of high pressure (Castañón et al. 2003; Gómez 2012; Hidalgo et al. 1998; Mendoza et al. 2002; Rodriguez and Preciado 1988; Scott and Rosenkranz 1977). However, the rare incidence of corn stunt disease in Argentina hinders the effective identification of resistant genotypes to this disease. The accurate detection of resistant genotypes may play a key role in the management of corn stunt disease in areas where the planting of temperate genotypes (obtained in areas where the prevalence of corn stunt is low or null) has increased over the past years ahead of tropical genotypes, which should *a priori* be more resistant to corn stunt for being obtained in a disease endemic area.

In the absence of high and constant natural pressures, the analysis of resistance mechanisms to corn stunt may be effective to select resistant genotypes (Azzam and Chancellor 2002; Carpane 2007), with the possibility to identify and combine several resistance mechanisms. For instance, the resistance of most genotypes is targeted to vector insects via antixenosis or antibiosis in other pathosystems (Rezaul Karim and Saxena 1991; Saxena 1987), thereby reducing the effectiveness of pathogen inoculation. In other cases, resistance seems to be aimed to the pathogen, since maize genotypes resistant to corn stunt show milder symptoms and a smaller yield reduction (Caro *et al*. 2008; 2009; Hidalgo *et al*. 1998).

The goal of this research was to identify the presence and to characterize resistance mechanisms to corn stunt disease in maize hybrids from temperate and tropical regions of Argentina, which would later allow to generate a screening methodology to be used in the absence of high corn stunt pressures.

## Materials and Methods

The tests implemented investigated the existence of different resistance mechanisms to corn stunt disease: antixenosis and antibiosis to vector *D. maidis* (Saxena 1987) and resistance to pathogen *S. kunkelii* (Hogenboom 1993).

### Biological Materials

A colony of healthy *D. maidis* was initiated from insects collected in the province of Tucumán (located in the tropical area of Argentina) and was maintained on plants of sweet corn variety Maizón at the IPAVE-CIAP (Plant Pathology Research Institute - Center of Agricultural Research (IPAVE-CIAP) at INTA (National Institute of Agricultural Technology) Córdoba, Argentina and at CEBIO (BioResearch Center) at UNNOBA-CICBA (National University of the North West of the Province of Buenos Aires - Scientific Research Commission of the Province of Buenos Aires), Pergamino, Buenos Aires, Argentina. The colonies were kept in aluminum-framed cages with a “*voile*” type nylon mesh, placed in a growth chamber at a temperature of 25°C, with a photoperiod of 16:8 (light: darkness) hours (Nault 1980).

Maize hybrids DK670 and DK72-10 which were selected in the temperate region of Argentina, and DK79-10 and DKB390 which were obtained from the tropical region of Argentina. The seed was obtained directly from Bayer’s seed processing facilities before they were treated, and for this reason they had no insecticides nor fungicides.

### Preference Test (Antixenosis to the vector *Dalbulus maidis*)

#### Test Conditions

The test was performed under controlled conditions at CEBIO at a mean temperature of 20-30°C using seeds planted in pots (1 seed per pot).

Plants with two fully expanded leaves were used. A plant from each hybrid was transferred into a glass cage. Pots were placed horizontally on the cage floor, so plants could be mounted in such a way that they exposed only a single leaf with its abaxial side upwards. Six two-week-old adult *D. maidis* mated females were later introduced into the cage.

Preference was determined by recording the number of insects settled on each hybrid or not settled on any hybrid at 1, 6, 24 and 48 hours (timepoints) after their release. At the end of the test, leaves were dissected using a binocular microscope to count the number of eggs.

#### Experimental Design and Statistical Analysis

The experiment was replicated 50 times, with each cage of four hybrids and six females serving as a replication. The arrangement of hybrids was randomized across replications. For the statistical analysis, a multinomial model was adjusted modeling hybrid and timepoints effects. The model was adjusted using the nnet package (Venables and Ripley 2002) of R language (R Core Team 2018). The response variable was the proportion of insects settled on each hybrid or on the cage. The significance of the differences in intercepts and slopes in the interaction was analyzed with contrasts using this same module. A generalized linear mixed model was used for oviposition to model negative binomial variables, with hybrid as fixed effect and replication as random effect.

### Survival Test (Antibiosis to the vector *Dalbulus maidis*)

#### Test Conditions

The test was performed under controlled conditions in IPAVE-CIAP at a temperature of 20-30°C using seeds planted in pots (1 seed per pot).

The test was conducted in plants with two fully expanded leaves. Each replication consisted of four polyethylene cages each containing a plant from one of the hybrids. Five two-week-old adult *D. maidis* mated females were then released. The number of surviving insects was counted weekly for four weeks (timepoints), and plants were replaced to ensure the presence of fresh plants.

#### Experimental Design and Statistical Analysis

The experiment was replicated 30 times, being a replication each group of four polyethylene cages containing each cage one hybrid and five insects. The distribution of cages was randomized across replication. For the statistical analysis, a generalized linear mixed model was used, being hybrid, timepoint and interaction fixed effects. The response variable was the probability of survival, with binomial distribution and a logit linkage function.

### Symptom Progression Test (Resistance to the pathogen *Spiroplasma kunkelii*)

To prevent reduced inoculation efficiency of pathogen *S. kunkelii* due to the potential presence of antixenosis and antibiosis to vector *D. maidis* by the hybrids tested, the artificial infestation took place using a high pressure of inoculative insects in no-choice conditions (Alivizatos and Markham 1986; Azzam and Chancellor 2002). This was considered enough to ensure that *S. kunkelii* was indeed inoculated to all plants as in other pathosystems (Shibata et al. 2007).

#### Test Conditions

A population of inoculative *D. maidis* was initiated by collecting symptomatic plants in Las Breñas, Province of Chaco, which were taken to IPAVE-CIAP. The acquisition of *S. kunkelii* from these plants and its subsequent maintenance was conducted according to Nault (1980) and adapted by Carpane (2007), in terms of acquisition access (7 days), incubation (21 days) and inoculation access (7 days) periods. Before the test, inoculative *D. maidis* were placed in glass test tubes (height: 15 cm; diameter: 2 cm) in groups of six adult insects and were transferred into the field where inoculation took place. Insects were not sexed, because transmission efficiency is similar in both genders (Alivizatos and Markham 1986).

Inoculation was performed in a field close to Monte Cristo, Province of Córdoba, at 30 km from IPAVE-CIAP. The presence of corn stunt disease in this area is typically low or null, thus minimizing the risk of natural infections from *D. maidis* interfering with forced inoculation. Plots of nine 500-meter-long rows were planted of each hybrid, with a row spacing of 52 cm and a density of 3.5 seeds per linear meter. At the four-leaf stage, five homogeneous blocks were labeled, and five plants of similar size were selected in each of them and individually covered with a “*voile*” type nylon cage. Insects were released into each cage (a tube containing six insects per plant) for an inoculation access period (IAP) of 48 hours, long enough to obtain maximum inoculation efficiency of *S. kunkelii* (Alivizatos and Markham 1986). As negative control, five plants of each hybrid were exposed to insects from the healthy colony (six adults per plant). Following the IAP, insects were controlled with insecticides and cages were removed. Insecticides were periodically sprayed later to prevent natural inoculation of *S. kunkelii* by eventual populations of *D. maidis* or by insects hatching from eggs laid in inoculated plants.

#### Progression of Disease Incidence and Severity

Symptom severity was assessed at 20, 45, 65 and 85 days after inoculation (DAI) timepoints. A 0-4 grade scale (based on Carpane *et al*. 2006) was used for the assessment, where 0 = no symptoms, 1 = leaves with red margins (red leaves), 2 = leaves with chlorotic stripes, 3 = leaves with mild stunting (height 15-30% of non-inoculated plants), 4 = leaves with severe stunting, with a height lower than 30% of non-inoculated plants.

#### Detection of the Pathogen *S*. *kunkelii*

In the last timepoint of symptom assessment (85 DAI), tissue samples from the ear leaf and the penultimate leaf from the tassel were collected to diagnose the presence of *S. kunkelii* at IPAVE-CIAP. A sample of 0.5 g leaf was cut and macerated in 5 mL of PBST (Shibata *et al*. 2007). The accumulation of *S. kunkelii* was estimated using a double antibody sandwich enzyme-linked immunosorbent assay (DAS-ELISA) conjugated with alkaline phosphatase (Giménez Pecci et al. 2009). *S. kunkelii* accumulation was expressed as relative absorbance (RA), this being the ratio between absolute absorbance at 405 nm of each leaf sample and a threshold of absorbance (mean + 3 standard deviations of absolute absorbance) obtained from six healthy plants of each hybrid (Sutula et al. 1986). Plants were considered positive (with presence of the pathogen *S. kunkelii*) if RA was higher to 1 in any of the leaves tested.

#### Determination of Yield

When grain moisture of DK72-10 (intermediate relative maturity of the hybrids tested) reached 16%, ears from each plant were harvested individually to determine yield (g/plant), which was expressed as qq/Ha, with an estimated planting density of 70,000 plants/Ha and a grain moisture of 14.5%. In turn, to neutralize the effect of yield potential *per se* of the hybrids tested, relative yield was calculated as the ratio between the yield of each plant and the average yield of ten non-inoculated plants from the same hybrid (all these plants were diagnosed negative for *S. kunkelii*) and expressed as percentage.

#### Experimental Design and Statistical Analysis

The statistical analysis of symptom progression was performed using a generalized mixed linear model with hybrid, timepoints (DAIs) and interaction as fixed effects. The response variables were binary (logit linkage): the incidence of plants with symptoms and plants with severe symptoms (grades of 3 or 4 in the scale mentioned above). The number of plants with a positive diagnose in DAS-ELISA was analyzed as the symptom progression considering only hybrid as fixed effect since samples were taken only once for diagnosis, and the incidence of plants with positive diagnosis in the ear leaf and the penultimate leaf as response variables. Accumulated severity (AS) of each plant was calculated by summing the severity grades across timepoints (DAIs). AS was analyzed in the same way as the number of plants with positive diagnose in DAS-ELISA. In turn, to analyze the relation between progression of symptom severity and RA, accumulated severity (AS) The relation between both variables was analyzed with Spearman’s rank correlation.

Yield was analyzed using a mixed linear model, with hybrid as fixed effect and block as random effect. Assumptions were validated through graphical analyses (residuals vs. predicted, normal QQ plot). Response variables were yield (qq/Ha) and relative yield (%). The relation between yield and AS was analyzed with mixed linear models, with hybrid, AS and their interaction as fixed effects. Linear models and generalized mixed models were adjusted using nlme (Pinheiro et al. 2018) and lme4 (Bates et al. 2015) packages of the R language (R Core Team 2018) through the statistical software interface InfoStat (Di Rienzo et al. 2018). Predicted values were compared using DGC test (Di Rienzo et al. 2002), with a significance level of 5% for all cases.

## Results

### Preference (Antixenosis to the vector *Dalbulus maidis*)

In the settling preference of adult *D. maidis* (Figure 1), a significant effect of hybrid and timepoint factors was observed, as well as their interaction (p<0.0001 in all cases). The hybrid effect was due to a higher number of insects settled on DK670 and DK79-10 than on DK72-10 and DKB390. The timepoint effect resulted from the increase in the number of insects settled on hybrids over time, as most insects settled on the cage rather than on hybrids at the first timepoints (mainly in the first hour of the test). The interaction between hybrid and timepoint (p=0.0213) was due to insects settling more rapidly over time in DK670 and DK79-10 than in DK72-10 and DKB390.

**Figure 1.**
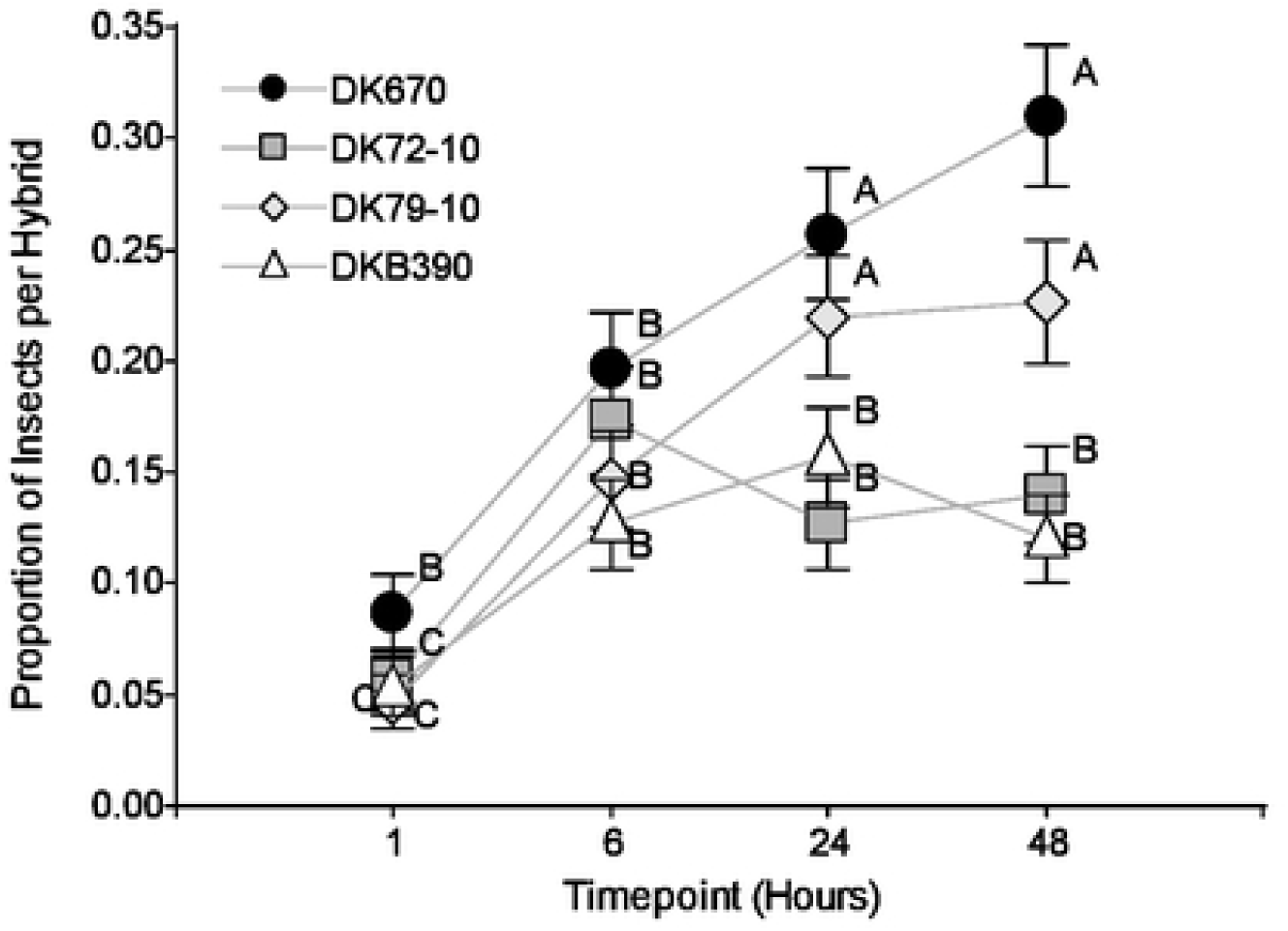
Proportion of *Dalbulus maidis* individuals settled on different hybrids overtime. Values sharing the same letter are not statistically different for a 5% significance level. Values with the same letter are not significantly different according to contrasts in the multinomial test (α= 0.05). Bars indicate standard error of the mean.

The number of eggs laid by *D. maidis* females on each hybrid (Figure 2) showed a significant effect of the hybrid factor (p<0.0001), with a similar arrangement to Figure 1, except that only DK72-10 had significantly fewer eggs than the other hybrids.

**Figure 2.**
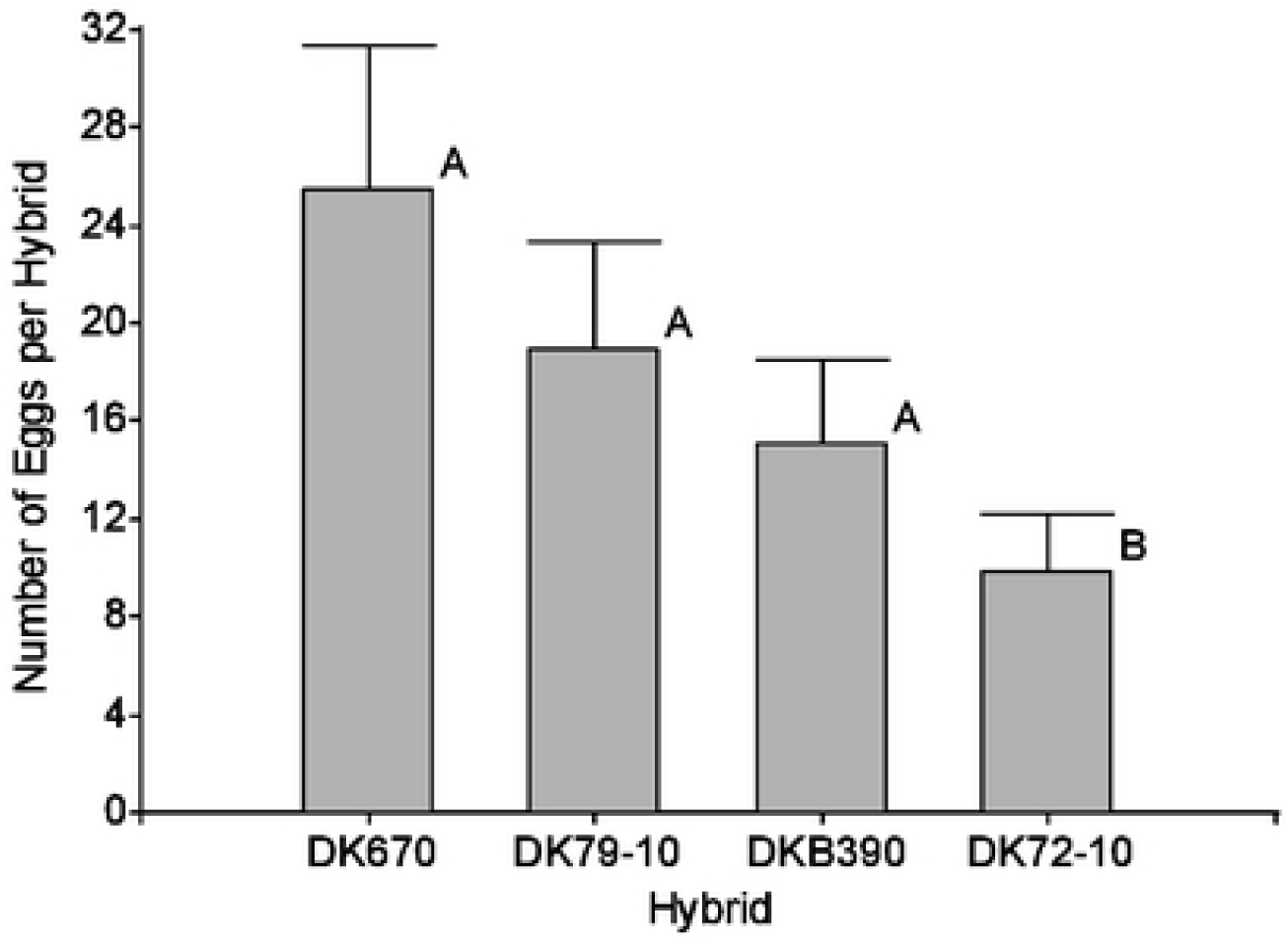
Number of eggs oviposited by *Dalbulus maidis* females during 48 hours in four maize hybrids. Values with the same letter are not significantly different according to contrasts in the mixed model test (α= 0.05). Bars indicate standard error of the mean.

### Survival (Antibiosis to the vector *Dalbulus maidis*)

For the probability of survival of *D. maidis* adults over time (Figure 3), a significant effect was seen for the hybrid and timepoint factors, as well as their interaction (p<0.0001 in all cases). The hybrid effect was due to the less survival in DK72-10 compared to other hybrids, and the timepoint effect resulted from the reduced survival over time in all hybrids. In turn, the interaction between hybrid and timepoint is explained by the faster decrease in survival in DK72-10 than in the other hybrids.

**Figure 3.**
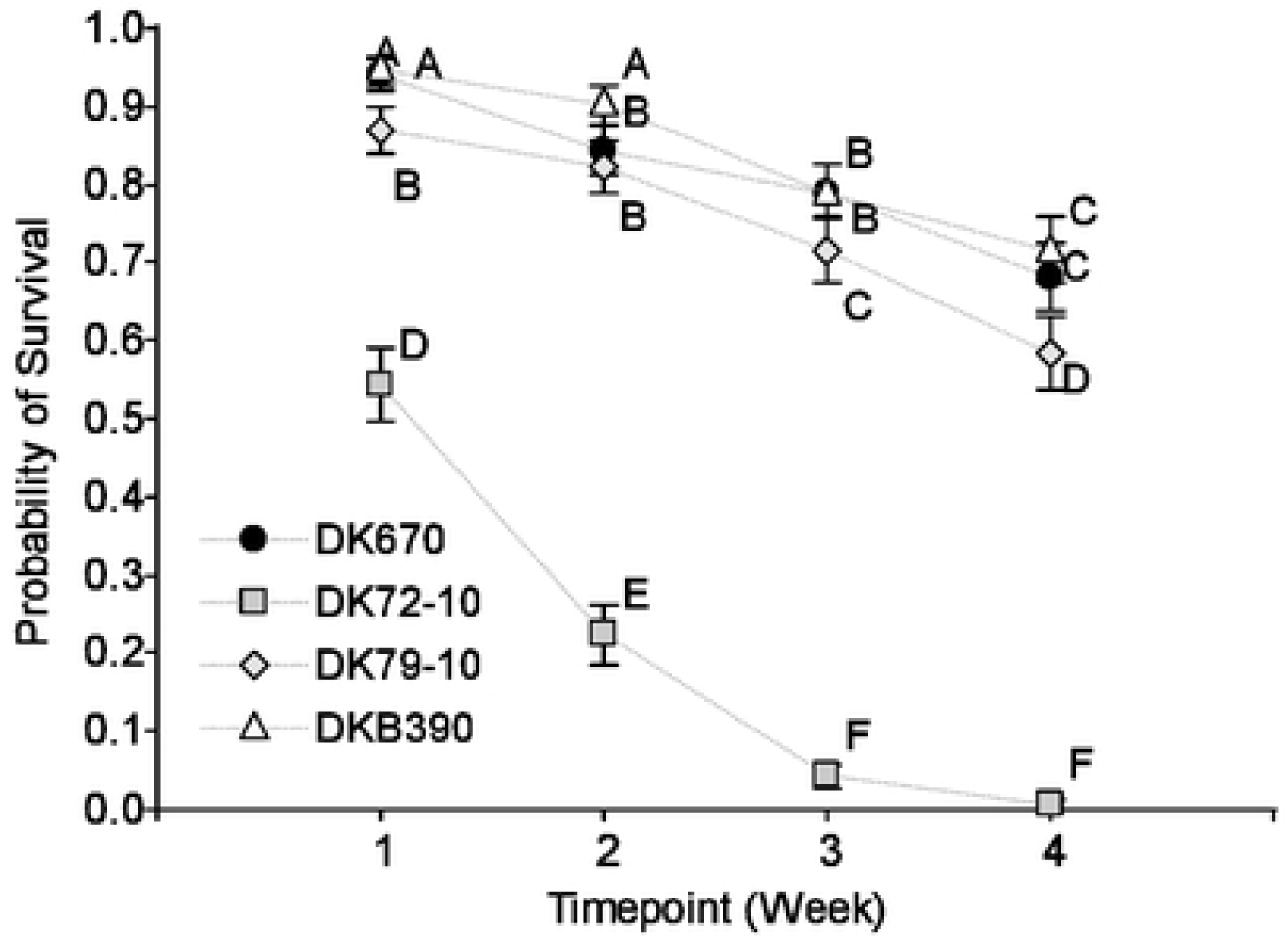
Probability of survival of *Dalbulus maidis* adults over time in four maize hybrids. Values sharing the same letter are not statistically different for a 5% significance level. Values with the same letter are not significantly different according to contrasts in the mixed model test (α= 0.05). Bars indicate standard error of the mean.

### Symptom Severity (Resistance to pathogen *Spiroplasma kunkelii*)

#### Progression of Disease Incidence and Severity

The incidence of plants with symptoms (Figure 4, left) showed a significant effect for hybrid (p<0.0001) and timepoint (p<0.0001) factors, as well as their interaction (p=0.0468). The hybrid effect was due to the incidence following the sequence DK670 = DK72-10 > DK79-10 > DKB390, the timepoint effect to the increase of incidence over time, and the interaction to a difference in the rate of increase of the proportion of plants with symptoms, following the order DK670 > DK72-10 > DK79-10 > DKB390. For instance, 96% of plants of DK670 and 68% of plants in DK72-10 showed symptoms at 45 DAI, while only 12% of plants of DK79-10 showed symptoms, and no symptomatic plants were found in DKB390 at this timepoint. Most plants developed symptoms after this period in these last two hybrids, with a higher rate in DK79-10 than in DKB390, which resulted in a higher final (at 85 DAI timepoint) incidence in the former hybrid than in the latter hybrid.

**Figure 4.**
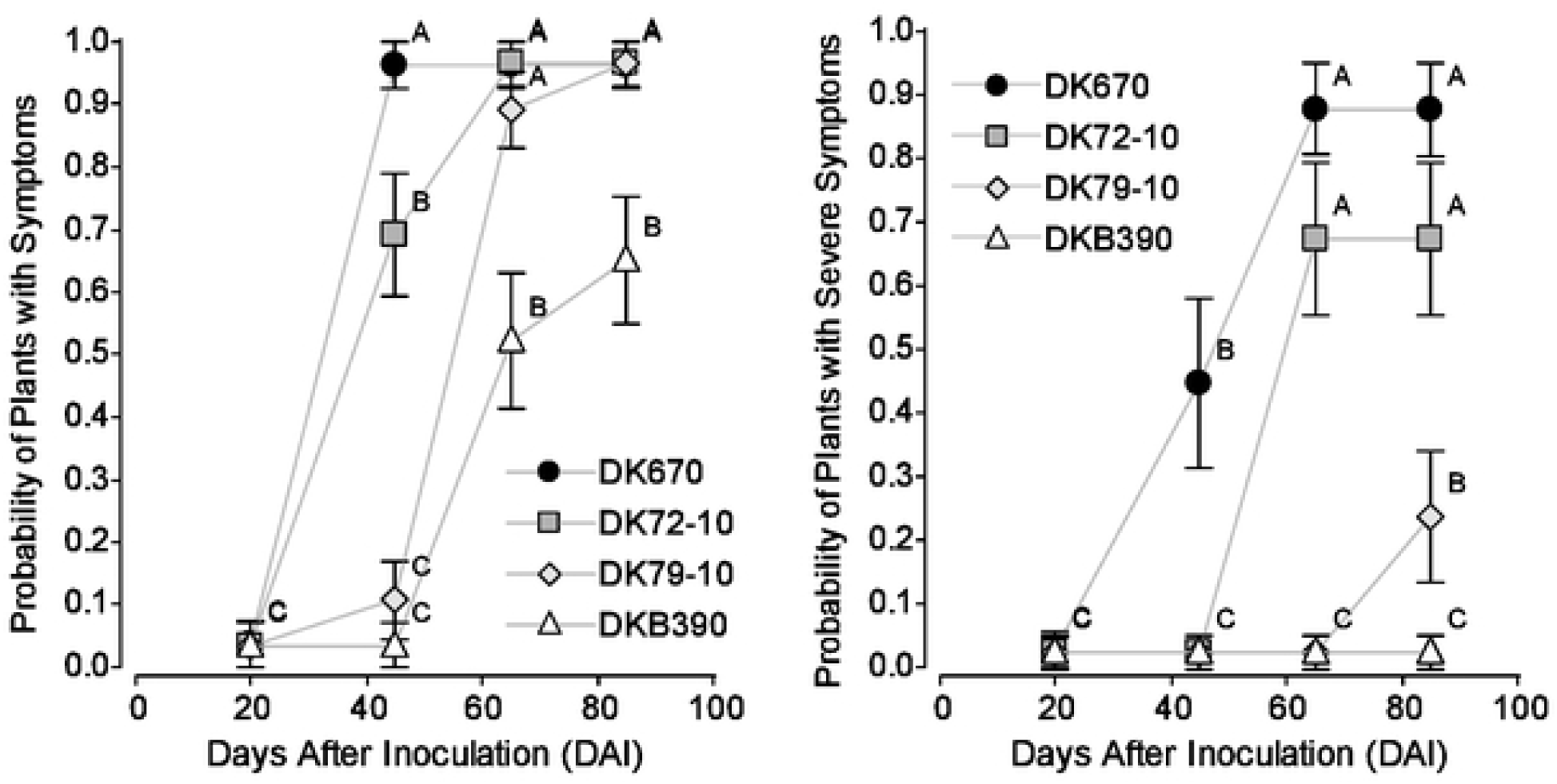
Incidence (Probability) of plants with symptoms (left) and with severe symptoms (right) after forced inoculation of the pathogen *Spiroplasma kunkelii* to four maize hybrids. Values sharing the same letter (within each panel) are not statistically different for a 5% significance level. Values with the same letter are not significantly different according to contrasts in the mixed model test (α= 0.05). Bars indicate standard error of the mean.

The incidence of plants with severe symptoms (Figure 4, right) showed a significant effect of the hybrid (p<0.0001) and timepoint (p<0.0001) factors, as well as their interaction (p=0.0483). The effect of the individual factors and their interaction was like that described for incidence of plants with mild + severe symptoms (Figure 4, left). At the end of the study, no DKB390 plants had severe symptoms

Figure 5 shows the sequence of symptom progression over time on a plant by plant basis. The hybrids tested had a similar sequence of symptoms, but with differences between them in the time of first detection and the following rate of progression. Symptoms started mostly as leaves with red margins or chlorotic stripes, followed by stunting. In some cases, symptoms were first seen as leaves with red margins followed by chlorotic stripes, although this sequence was less common. Finally, no remission of symptoms was seen in any case, i.e. plants showing symptoms at a certain timepoint kept displaying symptoms later, either similar or more severe.

**Figure 5.**
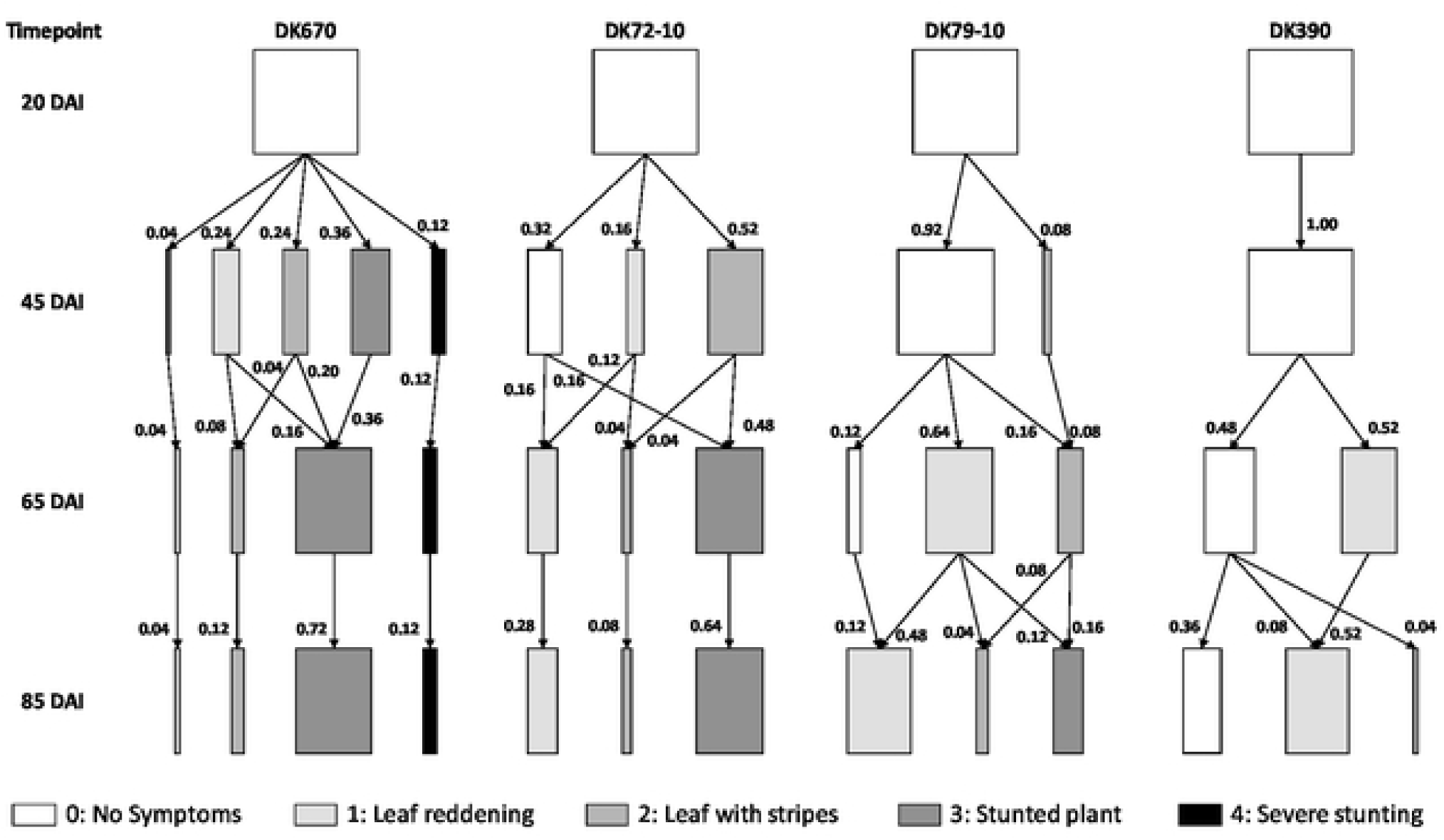
Kinematic diagram of the sequence of corn stunt disease symptom incidence and severity in four maize hybrids after forced inoculation at the four-leaf stage.

#### Detection of the Pathogen *S*. *kunkelii*

The comparison between the presence of corn stunt measured as symptoms and diagnosis using DAS-ELISA was performed on the penultimate (upper) leaf as it was somewhat more related with symptom severity than the ear leaf, mainly in plants that showed only mild symptoms after 65 DAI (Table 1). Four plants out of 100 tested negatives using the ear leaf and positive with the upper leaf (Table 1). Relative absorbances were almost 1 in the ear leaf in two plants (plant #9 in DK79-10 and #4 in DK390), and so the difference in diagnosis comparing the two leaves could be related to experimental error. However, there was a large difference in relative absorbance between both leaves for the other two plants, likely related to lack of detection of *S. kunkelii* in the ear leaf of these plants, which had displayed symptoms at the end of the test.

**Table 1.** Plants that differed in the result of the diagnosis for *Spiroplasma kunkelii* based on the ear leaf versus the upper leaf in DAS-ELISA tests, together with progression of symptom severity in these plants.

| Hybrid | Plant | Relative Absorbance (RA)<sup>#</sup> |  | Symptom Severity (at DAI) |  |  |  |
| --- | --- | --- | --- | --- | --- | --- | --- |
|  |  | Ear leaf | Upper leaf | 20 | 45 | 65 | 85 |
| DK79-10 | 9 | 0.903 | 1.561 | 0 | 0 | 0 | 1 |
|  | 11 | 0.375 | 3.005 | 0 | 0 | 1 | 3 |
| DKB390 | 4 | 0.935 | 1.114 | 0 | 0 | 0 | 0 |
|  | 19 | 0.126 | 1.703 | 0 | 0 | 0 | 1 |

<sup>#</sup> RA values below 1 indicate negative diagnosis (*S. kunkelii* not found), and above 1 positive diagnosis (presence of *S. kunkelii*).

All plants of DK670 and DK72-10 had symptoms and positive diagnosis for *S. kunkelii* (Table 2). In DK79-10, 92% of plants had symptoms and positive diagnosis, while 8% of plants with symptoms were diagnosed negative. These plants showed the first symptoms at 85 DAI as reddening of leaf margins (the mildest symptoms). All plants with symptoms in DKB390 (64%) tested positive. In addition, 20% of plants had positive diagnosis but no visible symptoms. Relative absorbances (RA) of these plants were low in the penultimate leaf (average 1.5) and lower to 1 in the ear leaf (that would have tested negative if only this leaf had been used for diagnosis). The remaining 16% of plants of this hybrid had no symptoms and tested negative for *S. kunkelii*.

**Table 2.**
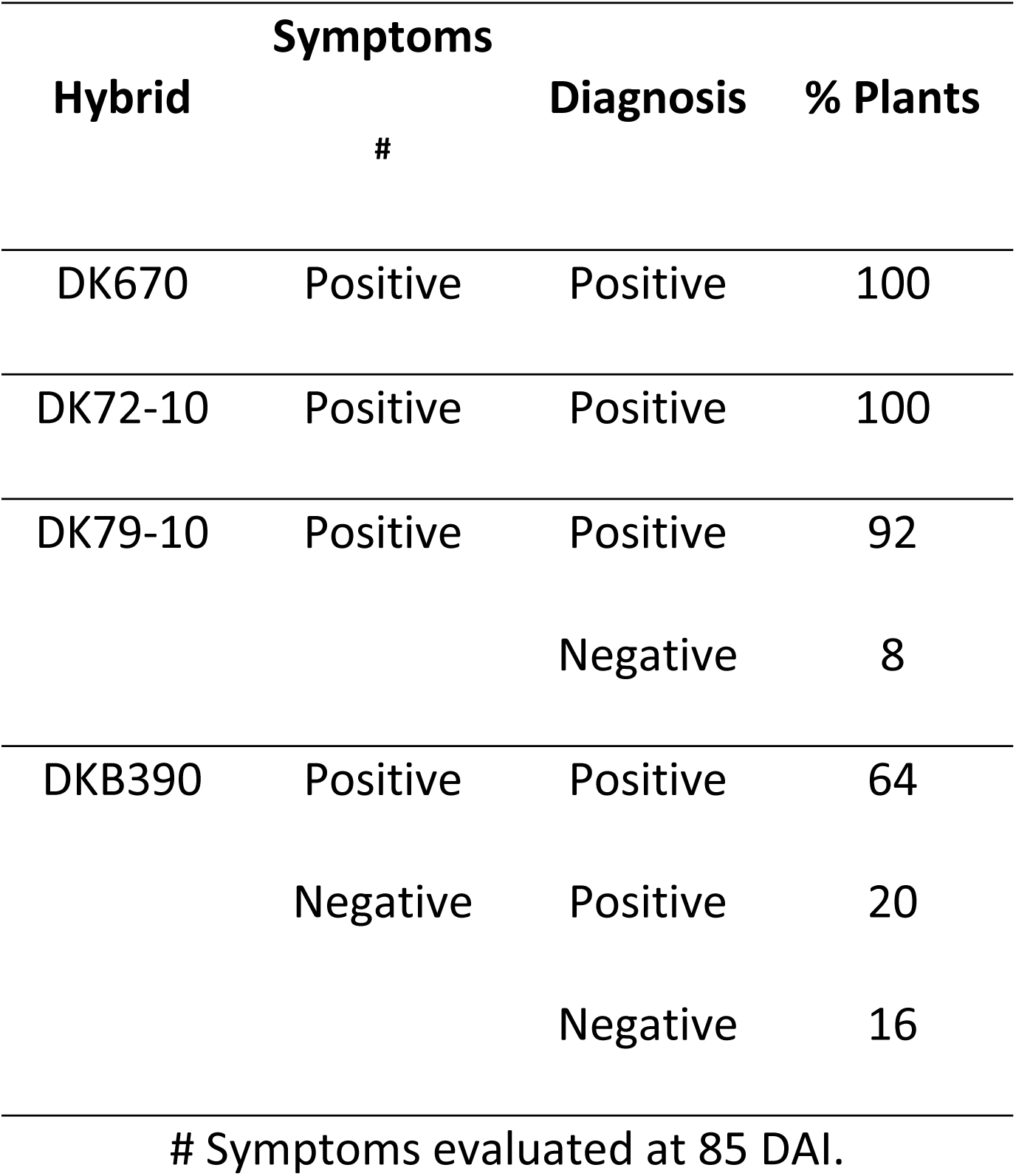
Proportion of plants with corn stunt symptoms and diagnosis for *Spiroplasma kunkelii* through DAS-ELISA after forced inoculation at the four-leaf stage in four maize hybrids.

The relation between AS and RA (Table 3) revealed a low correlation (no association) for all hybrids except DKB390. A positive and significant correlation was observed in this hybrid, thereby showing that more severe symptoms were associated with a greater accumulation of *S. kunkelii*.

**Table 3.** Spearman’s rank correlation between corn stunt accumulated severity (AS) and the accumulation of *Spiroplasma kunkelii* estimated as relative absorbance (RA) using DAS-ELISA in four maize hybrids.

| Hybrid | n | r Spearman | P value# |
| --- | --- | --- | --- |
| DK670 | 24 | 0.0617 | 0.7747 |
| DK72-10 | 25 | -0.0739 | 0.7254 |
| DK79-10 | 25 | -0.0090 | 0.9659 |
| DKB390 | 21 | 0.5070 | 0.0190 |

#### Determination of Yield

The yield of inoculated plants (Table 4) showed a significant effect for the hybrid factor (p<0.0001) and was directly correlated to incidence and severity of symptoms (Figure 4 and Figure 5) in the sequence DKB390 > DK79-10 > DK72-10 > DK670. The hybrid effect was also significant in terms of relative yield (p<0.0001), with a sequence DKB390 > DK79-10 > DK72-10 = DK670. The sequence of hybrids in both yield and relative yield is inversely related to accumulated severity; i.e. the hybrids with the highest yield had the lowest accumulated severity (AS), as shown in Table 4.

**Table 4.** Yield (qq/Ha), relative yield (RY, %) and accumulated severity (AS, rate) of four maize hybrids inoculated with *Spiroplasma kunkelii*.

| Hybrid | Yield (qq/Ha) |  | RY (%)<sup>#</sup> |  | AS (Rate)<sup>*</sup> |  |
| --- | --- | --- | --- | --- | --- | --- |
| DKB390 | 127.1 | A | 82.8 | A | 1.3 | D |
| DK79-10 | 92.4 | B | 50.2 | B | 3.0 | C |
| DK72-10 | 67.4 | C | 40.2 | C | 5.9 | B |
| DK670 | 46,2 | D | 37.1 | C | 8.0 | A |

<sup>#</sup> RY. Relative yield to non-inoculated plants.
<sup>\*</sup> AS. Sum of rates in four timepoints (maximum value of 16, as if severity rating was of 4 in every timepoint).
Values sharing the same letter (within each column) are not statistically different for a 5% significance level. Values with the same letter are not significantly different according to contrasts in the mixed model test ( $\alpha = 0.05$ ).

The effect of symptom severity on yield (Figure 6) was compared in two sections considering that AS was below 2 in DKB390. The first section analyzed the four hybrids in an AS from 0 to 2, and the second section compared the three remaining hybrids (excluding DKB390) in an AS from 3 to 7. In the first section, there was a significant effect for the hybrid (p=0.0013) and AS (<0.0001) factors, as well as their interaction (p=0.0008). The hybrid effect was due to the higher yield of DKB390, the AS effect to decreased yield resulting from the increase in AS, and the interaction to DKB390 having a higher yield than the others in AS of 0-1, without significant differences in an AS of 2. The coefficient of AS factor was -9.8, so was lowered by 9.8 qq/Ha for each unit of increase in AS. In the section of AS 3-7, only AS had a significant effect (p<0.0001), but not the hybrid (p=0.4345) nor the interaction between both factors (p=0.3428). The coefficient of the AS factor was -9.3 for this case, indicating that for each unit of increase in AS yield was reduced by 9.3 qq/Ha.

**Figure 6.**
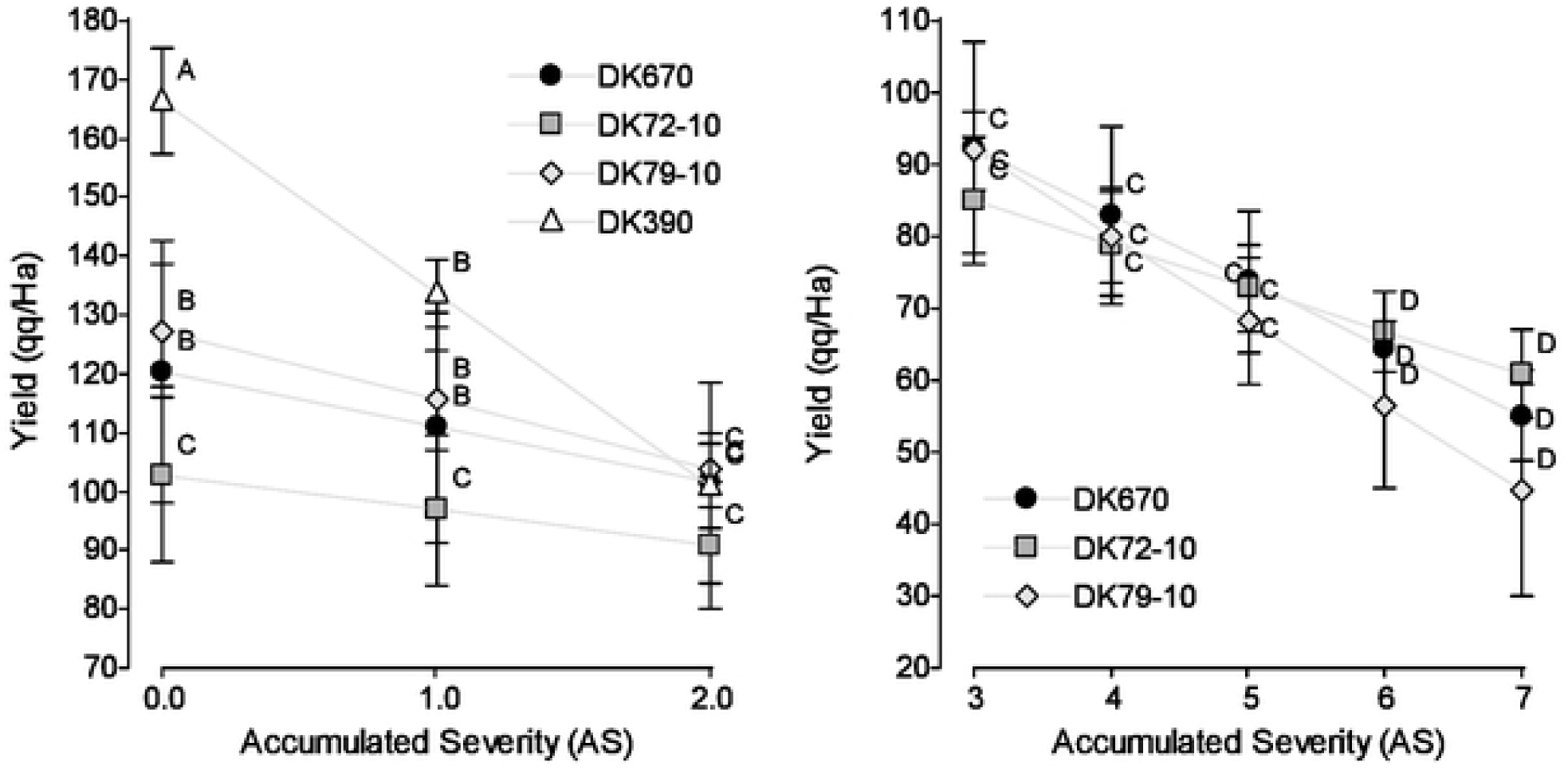
Relation between yield (qq/Ha) and accumulated severity (AS) of corn stunt disease symptoms. Left: Section with AS 0-2 in four maize hybrids. Right: Section with AS 3-7 excluding DKB390, because AS values were lower than 2. Values with the same letter within each panel are not significantly different according to contrasts in the mixed model test (α= 0.05). Bars indicate standard error of the mean.

## Discussion

Findings from this research prove that hybrids differ in both their level of resistance to corn stunt as well as the target organisms (either the insect vector or the pathogen) and the mechanisms of such resistance. Differences among hybrids were found in the level of antixenosis to *D. maidis*, resulting in decreased settling preference of *D. maidis* females, like findings in rice with *Nephotettix virescens* (Cheng and Pathak 1972; Heinrichs and Rapusas 1983a) and *Sogatella furcifera* (Khan and Saxena 1985; Ye and Saxena 1990), and in oats with *Delphacodes kuscheli* (Costamagna et al. 2005), where settling preference was also seen as early as six hours from exposure to plants. Our results suggest that maize hybrids DK72-10 and DKB390 negatively interfered with host acceptance by vector *D. maidis*, and hence may decrease the inoculation efficiency of the pathogen *S. kunkelii* to them, as such efficiency increases with the interval of insect-plant interaction (Alivizatos and Markham 1986). Settling preference tests using inoculative insects may confirm or rule out this hypothesis, which may be a future line of research.

Only DK72-10 showed antibiosis by reducing survival of *D. maidis* relative to other hybrids. An effect of antibiosis was also found in rice against *N. virescens* (Cheng and Pathak 1972; Heinrichs and Rapusas 1983a), *N. cincticeps* (Kawabe 1985), *Nilaparvata lugens* (Sogawa and Pathak 1970; Ye and Saxena 1990) and *S. furcifera* (Heinrichs and Rapusas 1983b; Khan and Saxena 1985). However, the antibiosis against *D. maidis* seen in this study was lower in absolute terms compared to other cases. For example, survival of *N. virescens* adults (Cheng and Pathak 1972; Ye and Saxena 1990) and *S. furcifera* (Heinrichs and Rapusas 1983b) was 20% in five days, while 58% of insects exposed to DK72-10 (the only one having antibiosis) survived one week. Hence, given that survival surpassed the 48-hour period in which maximum inoculation efficiency is achieved (Alivizatos and Markham 1986), antibiosis of DK72-10 would not decrease inoculation rate of *S. kunkelii*, although this antibiosis effect of maize against *D. maidis* may be present in other genotypes not tested here.

Regarding symptom progression in forced inoculation tests, we analyzed first inoculation efficiency among hybrids and how it was best diagnosed. In this sense, there was a highly close relation between diagnosis by symptom determination and DAS-ELISA, with a higher correlation when using the closest leaf to the tassel for the latter test. This is consistent with reports from Gussie et al. (1995) and Oliveira et al. (2002), who stressed that the probability of detection of *S. kunkelii* was higher in the apex of the plant, which may be explained by the fact that *S. kunkelii* moves with photosynthates to the actively growing apical regions of the plant, or that these parts promote the multiplication of *S. kunkelii*. In DK670 and DK72-10, all inoculated plants had visible symptoms and tested positive to *S. kunkelii*, but some DK79-10 plants that showed their first symptoms in the last timepoint were tested negatives. This may be due to the irregular distribution of *S. kunkelii* within the various plant organs and at low concentrations while displaying the first symptoms, thereby resulting in a negative diagnosis (Gussie et al. 1995).

Most plants in DKB390 (64%) showed symptoms and tested positive for *S. kunkelii*, but 20 % of plants showed no symptoms despite a positive diagnosis, and 16% of plants had no symptoms and a tested negative. On one hand, the presence of plants without symptoms but with positive diagnosis agrees with reports from Gussie et al. (1995), in which *S. kunkelii* is detected 10-30 days before symptom onset. Here, these plants might have shown symptoms in a later timepoint, although the test was completed when all hybrids were at physiological maturity. On the other hand, the presence of plants without symptoms and negative diagnosis may be due to: a) a failure in methodology, which seems unlikely because inoculation efficiency was 100% in other hybrids; b) antibiosis and/or antixenosis to *D. maidis*, lowering the length of insect-plant interaction and hence reducing inoculation efficiency of *S. kunkelii*, although DK72-10 had similar antixenosis and more antibiosis than DKB390 and all of its plants were inoculated, and in similar pathosystems (Hao and Pitre 1970; Shibata et al. 2007) these effects were neutralized using a load of inoculative insects similar to that used in this study; c) resistance of this hybrid to *S. kunkelii*, which may reduce movement or replication of this pathogen in the plant following inoculation, leading to *S. kunkelii* levels below the limit of detection of the method used (Gussie et al. 1995). In support of this latter motive, similar results were found for *S. kunkelii* in maize (Hidalgo et al. 1998; Massola Junior et al. 1999a; Oliveira et al. 2002) and for RTBV and RTSV in rice (Azzam and Chancellor 2002; Hibino et al. 1987), whom suggest that the target of resistance of some plant genotypes is the pathogen itself rather than the insect vector.

Hybrids differed in symptom progression, including initial detection, subsequent progress and final severity. Symptoms appeared earlier and progressed faster in susceptible vs. resistant hybrids as reported before (Massola Junior et al. 1999a; Scott et al. 1977). Based on these characters, the order of resistance to *S. kunkelii* was defined as DKB390 > DK79-10 > DK72 -10 > DK670. Hybrids from temperate areas, DK670 and DK72-10, were highly susceptible to *S. kunkelii*, as symptoms developed early (although later in DK72-10) and progressed rapidly reaching a high proportion of plants with severe damage and low yield. Symptoms in DK79-10 appeared later and progressed slowly, leading to a lower rate of plants with severe symptoms and intermediate yield. Finally, DKB390 was the most resistant hybrid to *S. kunkelii*, as some plants had no symptoms and no pathogen was found in them, others had no symptoms despite the presence of *S. kunkelii*, and others developed late-onset mild symptoms, resulting in the highest yield. These results confirm that higher yield reductions occur when symptoms appear early and progress rapidly as discussed before (Hidalgo et al. 1998; Massola Junior et al. 1999b; Oliveira et al. 2002; Scott et al. 1977), leading to two major concepts related to the control of this disease: a) the need to control corn stunt at early growth stages, way before symptom appearance, b) to identify resistant genotypes, rating symptom progression may be more useful than a single assessment at a specific timepoint. In this sense, an aspect to consider is that the lack of a complete correlation between symptom presence and diagnosis suggests that both are necessary to properly characterize genotypes resistant to corn stunt disease.

Yield loss caused by *S. kunkelii* ranging from 20% to 63% with final incidence rates between 63 and 100% are within the range previously reported (Hao and Pitre 1970; Hidalgo et al. 1998; Hruska and Gomez Peralta 1997; Massola Junior et al. 1999b; Scott et al. 1977; Virla et al. 2004). These authors also discussed that genotypes with similar levels of incidence and severity of symptoms can achieve different yields (Hidalgo et al. 1998; Oliveira et al. 2002; Scott et al. 1977;). This has been described as host tolerance, or the ability to obtain yield despite the damage caused by corn stunt. However, this response was not seen in this study, as yield reductions were directly related to the incidence and severity of symptoms.

No consistent relation was found between the accumulation of *S. kunkelii* and symptom severity, both intra- and inter hybrids. This contradicts other results (Caro et al. 2008; Gussie et al. 1995; Virla et al. 2004) reporting a higher accumulation in plants with severe symptoms but is consistent with findings in rice (Shibata et al. 2007), where no differences were observed in the accumulation of viruses causing Rice Tungro Disease among varieties differing in resistance or among plants with varying severity. The absence of a correlation in this work may be due to the method used (DAS-ELISA), which was performed here without a dilution curve to estimate the concentration of *S. kunkelii*. In turn, new methods to estimate pathogen accumulation (Okuda et al. 2019) found this relation for viruses causing Rice Tungro Disease, so therefore may be useful to characterize the dynamics of *S. kunkelii* accumulation in maize hybrids differing in resistance to corn stunt.

This work identified resistance mechanisms to the vector *D. maidis* and to the pathogen *S. kunkelii* as components of corn stunt pathosystem. Hybrids differed in the level of resistance and target organisms of such resistance. DK72-10 showed antixenosis and antibiosis to *D. maidis*, DK79-10 was resistant to *S. kunkelii* and DKB390 expressed antixenosis to *D. maidis* and resistance to *S. kunkelii*. More than one mechanism and target organism (vector or pathogen) of resistance were identified with this strategy, which therefore provides the potential to combine them to obtain even more resistant hybrids. This is an advantage of this type of testing over natural infestations, where the observed response (incidence or severity of symptomatic plants) does not allow to differentiate between these foregoing factors. In summary, the presence of resistant hybrids combined with the use of an effective method for differentiation may facilitate the selection of hybrids capable of reducing the negative impact of corn stunt.

## Acknowledgements

We thank Mariana Ferrer for his assistance in setting up insect survival and symptom progression experiments.

This research did not receive any specific grant from funding agencies in the public, commercial, or not-for-profit sectors.

## References

Alstatt, G. 1945. A new corn disease in the Rio Grande Valley. Plant Disease. Rep. 29: 533–534.

Alivizatos, A.S., and Markham, P.G. 1986. Acquisition and transmission of corn stunt spiroplasma by its leafhopper vector *Dalbulus maidis*. Ann. Appl. Biol. 108: 535–544.

Azzam, O., and Chancellor, T. 2002. The Biology, Epidemiology, and Management of Rice Tungro Disease in Asia. Plant Disease 86: 88–100.

Bates, D., Maechler, M., Bolker, B., and Walker, S. 2015. Fitting Linear Mixed-Effects Models Using lme4. Journal of Statistical Software 67: 1–48.

Bajet, N.B., and Renfro, B.L. 1989. Occurrence of corn stunt spiroplasma at different elevations in Mexico. Plant Disease 73: 926–930.

Bradfute, O.E., Tsai, J.H., and, Gordon D.T. 1981. Corn Stunt Spiroplasma and viruses associated with a maize disease in Southern Florida. Plant Disease 65: 837–841.

Carloni, E. 2010 Características de corn stunt spiroplasma en Argentina. M Sc. Tesis. Facultad de Ciencias Agropecuarias. Universidad Nacional de Córdoba.

Carloni, E., Carpane, P., Paradell, S., Laguna, I., and Giménez Pecci, M.P. 2013. Presence of *Dalbulus maidis* (Hemiptera: Cicadellidae) and of *Spiroplasma kunkelii* in the Temperate Region of Argentina. J. Econ. Entomol. 106: 1574–1581.

Carloni, E., Virla, E., Paradell, S., Carpane, P., Nome, C., Laguna, I., and Giménez Pecci, M.P. 2011. *Exitianus obscurinervis* (Hemiptera: Cicadellidae), a new experimental vector of *Spiroplasma kunkelii*. J. Econ. Entomol. 104: 1793–1799.

Caro, L.A., Giménez Pecci, M.P., and Laguna, I.G. 2008. Susceptibilidad de 3 genotipos de maíz a *D*. *maidis* y a *Spiroplasma kunkelii* evaluada mediante variables fisiológicas y de rendimiento. 1° Congreso Argentino de Fitopatología, Córdoba, 28-30 mayo. Resúmenes (ISBN 978-987-24373-0-51): 336.

Caro, L.A., Bisonard, E.M., Laguna, I.G., and Giménez Pecci, M.P. 2009. Efecto del corn stunt spiroplasma (CSS) sobre parámetros de rendimiento de tres genotipos de maíz. XIII Jornadas Fitosanitarias Argentinas, Termas de Río Hondo, Santiago del Estero, 30 Sept, 1 y 2 oct 2009.

Carpane, P., Laguna, I.G., Virla, E., Paradell, S., and Giménez Pecci, M.P. 2006. Experimental transmission of corn stunt spiroplasma present in different regions of Argentina. Maydica 51: 461–468.

Carpane, P. 2007. Host resistance and diversity of *Spiroplasma kunkelii* as components of corn stunt disease. PhD. Thesis. Department of Entomology and Plant Pathology. Oklahoma State University. Stillwater, OK., USA.

Carpane, P., Melcher, U., Wayadande, A., Giménez Pecci, M.P., Laguna, I.G., Dolezal, W., and Fletcher, J. 2013. An analysis of the genomic variability of the phytopathogenic mollicute *Spiroplasma kunkelii*. Phytopathology 103: 129–134.

Castañón, G., Hidalgo, H., and Jeffers, D. 2003. Heterosis en siete líneas de maíz para tolerancia al achaparramiento y rendimiento de grano.

Cheng, C.H., and Pathak, M.D. 1972. Resistance to *Nephotettix virescens* in rice varieties. J. Econ. Entomol. 65: 1148–1153.

Costamagna, A.C., Remes Lenicov, A.M.M., and Zanelli, M. 2005. Maize and oat antixenosis and antibiosis against *Delphacodes kuscheli* (Homoptera: Delphacidae), vector of “Mal de Rio Cuarto” of maize in Argentina. J. Econ. Entomol. 98: 1374–1381.

Di Rienzo, J.A., Guzmán, A.W., and Casanoves, F. 2002. A Multiple Comparisons Method Based on the Distribution of the Root Node Distance of a Binary Tree. J. Agric. Biol. Environ. Stat. 7:1–14.

Di Rienzo, J.A., Casanoves, F., Balzarini, M.G., Gonzalez, L., Tablada, M., and Robledo, C.W. InfoStat Version 2015. Grupo InfoStat, FCA, Universidad Nacional de Córdoba, Argentina. URL http://www.infostat.com.ar.

Frazier, N. 1945. A streak disease of corn in California. Plant Dis. Rep. 29: 212–213.

Giménez Pecci, M.P, Carpane, P., Nome, C., Paradell, S., Remes Lenicov, A., Virla, E., and Laguna, I.G. 2003. Presencia del CSS y su vector *Dalbulus maidis* en el noreste argentino. Fitopatol. Brasileira 28: 280.

Giménez Pecci, M.P., Laguna, I.G., Carpane, P.D., Carloni, E., and Murúa, L. 2005. Dispersión e incidencia del corn stunt spiroplasma en el cultivo de maíz en diferentes áreas de Argentina. XIII Congreso Latinoamericano de Fitopatología y III Taller de la Asociación Argentina de Fitopatólogos. 19-22 abril. Córdoba, Argentina Res 477.

Giménez Pecci, M.P., Laguna, I.G., Ávila, A.O., Remes Lenicov, A.M., Virla, E., Borgogno, C., Nome, C.F., and Paradell, S. 2002. Difusión del corn stunt spiroplasma del maíz (*Spiroplasma kunkelii*) y del vector (*Dalbulus maidis*) en la República Argentina. Revista de la Facultad de Agronomía de La Plata 105: 1–8.

Gómez, L.G. 2012. Cap. II. El maíz en áreas del trópico. Pg. 25–30. En: Enfermedades del maíz producidas por virus y mollicutes en Argentina. Editores: Giménez Pecci, MP, Laguna, IG, Lenardón SL. Ediciones INTA (ISBN: 978-987-679-116-8).

Gussie, J.S., Fletcher, J., and Claypool, P.L. 1995. Movement and multiplication of *Spiroplasma kunkelii* in corn. Phytopathology 85: 1093–1098.

Hao, G., and Pitre, H. 1970. Relationship of Vector Numbers and Age of Corn Plants at Inoculation to Severity of Corn Stunt Disease. Journal of Economic Entomology 63: 924–927.

Heinrichs, E.A., and Rapusas, H. 1983a. Correlation of resistance to the green leafhopper, *Nephotettix virescens* (Homoptera: Cicadellidae) with tungro virus infection in rice varieties having different genes for resistance. Environ. Entomol. 12: 201–205.

Heinrichs, E.A., and Rapusas, H. 1983b. Levels of resistance to the whitebacked planthopper, *Sogatella furcifera* (Homoptera: Delphacidae), in rice varieties with different resistance genes. Environ. Entomol. 12: 1793–1797.

Hibino, H., Tiongco, E.R., Cabunagan, R.C., and Flores, Z.M. 1987. Resistance to rice tungro-associated viruses in rice under experimental and natural conditions. Phytopathology 77: 871–875.

Hidalgo, H., Jeffers, D., and Rodríguez, F. 1998. Resistencia al achaparramiento del maíz mediante infestaciones de *Dalbulus maidis* en maíz. Agronomía Mesoamericana 9: 119–124.

Hogenboom, N.G. 1993. Economic importance of breeding for disease resistance, pp. 5–9. In T. Jacobs and J. Parlevliet [eds.], Durability of disease resistance. Kluwer Academic Publishers, Boston.

Hruska, A.J., Gladstone, S.M., and Obando, R. 1996. Epidemic roller coaster: maize stunt disease in Nicaragua. Am. Entomology 42: 248–252.

Hruska, A.J, and Gómez Peralta, M. 1997. Maize Response to Corn Leafhopper (Homoptera: Cicadellidae) Infestation and Achaparramiento Disease. J. Econ. Entomol. 90: 604–610.

Kawabe, S. 1985. Mechanism of varietal resistance to the rice green leafhopper (*Nephotettix cincticeps* Uhler). JARQ 19: 115–124.

Khan, Z. R., and Saxena, R.C. 1985. Behavioral and physiological responses of *Sogatella furcifera* (Homoptera: Delphacidae) to selected resistant and susceptible rice cultivars. J. Econ. Entomol. 78: 1280–1286.

Kunkel, L. 1946. Leafhopper transmission of corn stunt. PNAS 32: 246–247.

Lenardón, S.L., Laguna, I.G., Gordon, D.T., Truol, G.A., Gomez, J., and Bradfute, O.E. 1993. Identification of Corn Stunt Spiroplasma in maize from Argentina. Plant Disease 77: 100.

Massola Júnior, N.S., Bedendo, I.P., Amorim, L., and Lopes, J.R.S. 1999a. Fitoplasma e espiroplasma em milho: multiplicação e efeito na produção de genótipos resistente e suscetível. Summa Phytopathologica 25: 356–359.

Massola Júnior, N.S., Bedendo, I.P., Amorim, L., and Lopes, J.R.S. 1999b. Effects of the inoculation time on corn with *Spiroplasma kunkelii* on yield components. Fitopatologia Brasileira 24: 571–573.

Mendoza, M., López, A., Rodríguez, S., Oyervides, García A., De León, C., and Jeffers, D. 2002. Acción génica de la resistencia al achaparramiento del maíz causado por espiroplasma, fitoplasmas y virus. Revista Mexicana de Fitopatología 20: 13–17.

Nault, L.R. 1980. Maize bushy stunt and corn stunt: A comparison of disease symptoms, pathogen host ranges, and vectors. Phytopathology 70: 659–662.

Nault, L.R. 1990. Evolution of an insect pest: maize and the corn leafhopper, a case study. Maydica 35: 165–175.

Okuda, M., Shiba, T., Hirae, M., Masunaka, A, and Takeshita, M. 2019. Analysis of symptom development in relation to quantity of *Rice stripe virus* in rice (*Oryza sativa*) to simplify evaluation of resistance. Phytopathology 109: 701–707.

Oliveira, E., Waquil, J.M., Fernandes, F.T., Paiva, E., Resende, R., and Kitajima, W.E. 1998. Enfezamento pálido e enfezamento vermelho na cultura do milho no Brasil Central. Fitopatologia Brasileira 23: 45–47.

Oliveira, E., Oliveira, C., Souza, I., Magalhaes, P. and Cruz, I. 2002. Enfezamentos em milho: expressão de sintomas foliares, detecção dos molicutes e interações com genótipos. Revista Brasileira de Milho e Sorgo 1: 53–62.

Oliveira, E., Resende, R., Gimenez Pecci, M.P., Laguna, I.M., Herrera, P., and Cruz, I. 2003. Incidência de viroses e enfezamentos e estimativa de perdas causadas por molicutes em milho no Paraná. Pesquisa Agropecuária Brasileira 38: 19–25.

Paradell, S., Virla, E., and Toledo, A. 2001. Leafhoppers species richness and abundance on corn crops in Argentina (Insecta-Hemiptera-Cidadellidae). Bol. Sanidad Vegetal y Plagas 27: 465–474.

Pinheiro, J., Bates, D., DebRoy, S., Sarkar, D., and R Core Team. 2018. nlme: Linear and Nonlinear Mixed Effects Models. R package version 3.1-137. URL: https://CRAN.R-project.org/package=nlme.

R Core Team. 2018. R: A language and environment for statistical computing. R Foundation for Statistical Computing, Vienna, Austria. https://www.R-project.org/.

Rezaul Karim, A., and Saxena, R.C. 1991. Feeding behavior of three *Nephotettix* species (Homoptera: Cicadellidae) on selected resistant and susceptible rice cultivars, wild rice, and graminaceous weeds. J. Econ. Entomol. 84: 1208–1215.

Rodriguez, E.A., and Preciado, R.E. 1988. Obtención de variedades resistentes o tolerantes a enfermedades foliares y de mazorca. Primera Reunión Científica y Agropecuaria. Centro de Investigaciones Forestales y Agropecuarias de Veracruz. INIFAPCIRGOC. MEX. p. 17.

Saxena, R.C. 1987. Antifeedants in tropical pest management. Insect Sci. Application 8: 731–736.

Scott, G., and Rosenkranz, E. 1977. Effectiveness for Recurrent Selection for Corn Stunt Resistance in a Maize Variety. Crop Science 14: 758–760.

Scott, G., Rosenkranz, E., and Nelson, L. 1977. Yield losses of corn due to corn stunt disease complex. Agronomy Journal 69: 92–94.

Shibata, Y., Cabunagan, R.C., Cabauatan, P.Q., and Choi, I.R. 2007. Characterization of *Oryza rufipogon*–derived resistance to tungro disease in rice. Plant Disease 91: 1386–1391.

Sogawa, K., and Pathak, M.D. 1970. Mechanisms of brown planthopper resistance in Mudgo variety of rice (Hemiptera: Delphacidae). Appl. Entomol. Zool. 5: 145–158.

Sutula, C., Gillet, J., Morrisey, S., and Ramsdell, D. 1986. Interpreting ELISA data and establishing the positive-negative threshold. Plant Disease 70: 722–726.

Venables, W.N., and Ripley, B.D. 2002. Modern Applied Statistics with S. Fourth Edition. Springer, New York. ISBN 0-387-95457-0.

Virla, E., Remes Lenicov, A., and Paradell, S. 1990. Presencia de *Dalbulus maidis* sobre maíz y teosinte en la República Argentina (Insecta - Homoptera - Cicadellidae). Rev. Fac. de Agr. La Plata. 66/67: 23–30.

Virla, E., Paradell, S., and Diez, P. 2003. Estudios bioecológicos sobre la chicharrita del maíz *Dalbulus maidis* (Insecta - Cicadellidae) en Tucumán (Argentina). Bol. San. Veg. Plagas 29: 17–25.

Virla, E.G., Díaz, C.G., Carpane, P.D., Laguna, I.G., Ramallo, J., Gómez, G.L., and Giménez Pecci, M.P. 2004. Evaluación preliminar de la disminución en la producción de maíz causada por el “Corn stunt spiroplasma” (CSS) en Tucumán, Argentina. Bol. San. Veg. Plagas 30: 257–267.

Whitcomb, R.F., Chen, T.A., Williamson, D.L., Liao, C., Tully, J.G., Bove, J.M., Mouches, C., Rose, D.L., Coan, M.E., and Clark, T.B. 1986. *Spiroplasma kunkelii* sp. nov.: characterization of the etiological agent of corn stunt disease. J. Syst. Bacteriol. 36: 170–178.

Ye, Z.H., and Saxena, R.C. 1990. Resistance to Whitebacked Planthopper in elite lines of cultivated X wild rice crosses. Crop Sci. 30: 1178–1182.

